# Physics-Informed Operator Learning for Pulsatile Milk Flow in Distal Generations of a Bifurcated Mammary Duct Network

**DOI:** 10.64898/2026.06.12.731941

**Authors:** Abdullahi Olapojoye, Aria Nosratinia, Fatemeh Hassanipour

## Abstract

Pulsatile milk transport through the lactating mammary ductal tree involves complex interactions between pressure gradients, wall compliance, and non-Newtonian rheology across spatial scales that span nearly two orders of magnitude in lumen radius. Direct experimental characterisation of flow in distal ductal generations remains infeasible due to their sub-millimetre calibre, leaving the haemodynamic environment of the secretory ductules largely unknown. We present a two-stage physics-informed operator-learning framework that extends validated flow predictions from three instrumented duct generations to twenty generations of a bifurcated mammary network. A Physics-Informed Neural Network (PINN) trained against particle image velocimetry measurements across seven ducts achieved *R*^2^ = 0.924–0.997. A Deep Operator Network (DeepONet) distilled from the PINN and refined through physics-constrained training on the governing one-dimensional fluid–structure interaction equations achieved *R*^2^(*u*) = 0.857–0.985 across all validated ducts, with predictions for Generations 4–20 obtained by supplying Murray’s Law geometry and mass-conservation-scaled boundary conditions to the frozen operator. Three biophysically significant findings emerge: a mean velocity plateau of 0.14–0.18 m/s across Generations 4–13 produced by Cross shear-thinning compensation offsetting Murray-branching deceleration; a non-monotonic pulsatility index that declines from 0.048 at Generation 1 to a minimum of 0.039 at Generation 5 before rising monotonically to 1.37 at Generation 20 as progressive wall stiffening drives the most distal ductules into a microcirculation-like haemodynamic regime; and a brief elastic-recoil transition zone at Generations 4–5 where mean axial pressure drop reverses sign. To the authors’ knowledge, these results provide the first quantitative characterisation of pulsatile milk flow across the full hierarchy of a bifurcated mammary ductal tree using a physics-informed operator-learning framework with implications for ductal mechanobiology, milk ejection mechanics, and mastitis pathogenesis.

## 1 Introduction

The mammary ductal system forms a hierarchical bifurcating network in which a small number of collecting ducts emerge from the nipple and progressively branch into smaller conduits terminating in secretory alveoli [1]. Ultrasound imaging studies have shown that the lactating human breast contains between four and eighteen major ductal openings at the nipple, each draining an independent ductal tree [2, 3]. Duct lumen diameters decrease from approximately 1.5 mm in the proximal collecting ducts to tens of micrometres in the terminal ductules, producing a geometric hierarchy spanning nearly two orders of magnitude in length scale. During breastfeeding, oscillatory suction generated by infant suckling drives pulsatile milk transport through this compliant network. The resulting flow is governed by coupled interactions among intraductal pressure gradients, duct wall deformation, and the non-Newtonian rheology of human milk.

These fluid mechanical processes are directly relevant to both lactation physiology and breast health. Impaired ductal transport has been implicated in lactation mastitis, a painful inflammatory condition affecting up to one third of breastfeeding women [1]. In addition, the mechanical environment experienced by mammary epithelial cells including wall shear stress, transmural pressure, and cyclic lumen deformation, influences mechanobiological signaling within the secretory alveoli [4]. Quantitative characterization of intraductal flow is therefore important for understanding milk ejection, ductal patency, and epithelial cell response during lactation.

Mathematical modeling of milk transport in the lactating breast has remained comparatively limited. Early studies adopted simplified rigid-tube formulations with Newtonian flow assumptions [5]. Mortazavi et al. [6] later examined the fluid dynamics of milk ejection and demonstrated that intraductal pressure during breastfeeding is governed primarily by the milk ejection reflex rather than infant suction alone. More recently, Olapojoye et al. [7] combined computational fluid dynamics and machine learning to characterise pulsatile flow within a single proximal duct network. A subsequent study extended this framework to a compliant three-generation bifurcating duct network using domain-decomposed Physics-Informed Neural Networks (PINNs), incorporating the non-Newtonian rheology of human milk and validating predictions against particle image velocimetry (PIV) measurements obtained from a bio-inspired breastfeeding simulator [8, 9]. Despite these advances, existing computational studies remain restricted to the proximal duct generations accessible to imaging and instrumentation. Consequently, flow conditions within distal duct generations which are directly relevant to milk stasis and epithelial mechanobiology - remain largely uncharacterised.

The one-dimensional (1D) compliant-tube framework developed for arterial haemodynamics [10] provides a computationally efficient approach for modeling pulsatile flow in branching elastic networks. In this framework, cross-sectionally averaged conservation equations are coupled with a pressure–area constitutive relation to capture the dominant fluid–structure interaction and wave propagation behaviour at substantially lower computational cost than fully three-dimensional simulations [11, 12]. The approach has been extensively validated in arterial networks [13, 14] and pulmonary airways [15], which exhibit branching geometries and compliance distributions analogous to those of the mammary ductal tree. Furthermore, the Womersley number in mammary ducts satisfies Wo ≪ 1 throughout the network (Section 6.2), supporting the quasi-parabolic velocity profile assumption underlying the 1D friction model. Because imaging data for distal duct generations are unavailable, network geometry in the present study is extrapolated using Murray’s Law of minimum work [16], which predicts a symmetric daughter-to-parent radius ratio of 2^−1*/*3^ ≈ 0.794. This scaling relationship is consistent with proximal branching ratios measured experimentally by Ramsay et al. [2].

Alongside these developments in reduced-order modeling, physics-informed machine learning methods have emerged as promising tools for solving inverse and data-assimilation problems in biofluid mechanics. Physics-Informed Neural Networks (PINNs) [17] incorporate governing partial differential equations directly into the network loss function, enabling reconstruction of flow fields from sparse experimental measurements [18, 19]. However, PINNs typically require retraining for each new geometry or boundary condition, limiting their efficiency in hierarchical networks with many generations. This limitation becomes particularly significant in mammary duct networks, where duct dimensions and flow conditions vary systematically across successive bifurcations. Neural operators address this challenge by learning mappings between functional spaces rather than individual solutions [20, 21]. Among these approaches, the Deep Operator Network (DeepONet) architecture [21] has demonstrated strong capability for generalising flow solutions across unseen parameter regimes and geometries. Combining a validated PINN framework with a DeepONet surrogate therefore provides a pathway for extending experimentally constrained predictions beyond the limited set of measurable duct generations.

The present study addresses the lack of quantitative data in distal mammary ducts by developing a two-stage physics-informed operator-learning framework capable of extending flow predictions from experimentally instrumented proximal ducts to a twenty-generation bifurcating network. In the first stage, a PINN reconstructs spatiotemporal pressure, velocity, and lumen-area dynamics within a seven-duct compliant phantom by simultaneously solving the governing 1D fluid–structure interaction equations and assimilating PIV velocity measurements from nine sensor locations [9]. In the second stage, a DeepONet trained on physics-consistent PINN predictions generalises the learned dynamics to unseen duct generations and boundary conditions. The bifurcating network topology considered in this study is illustrated in Figure 1.

**Figure 1.**
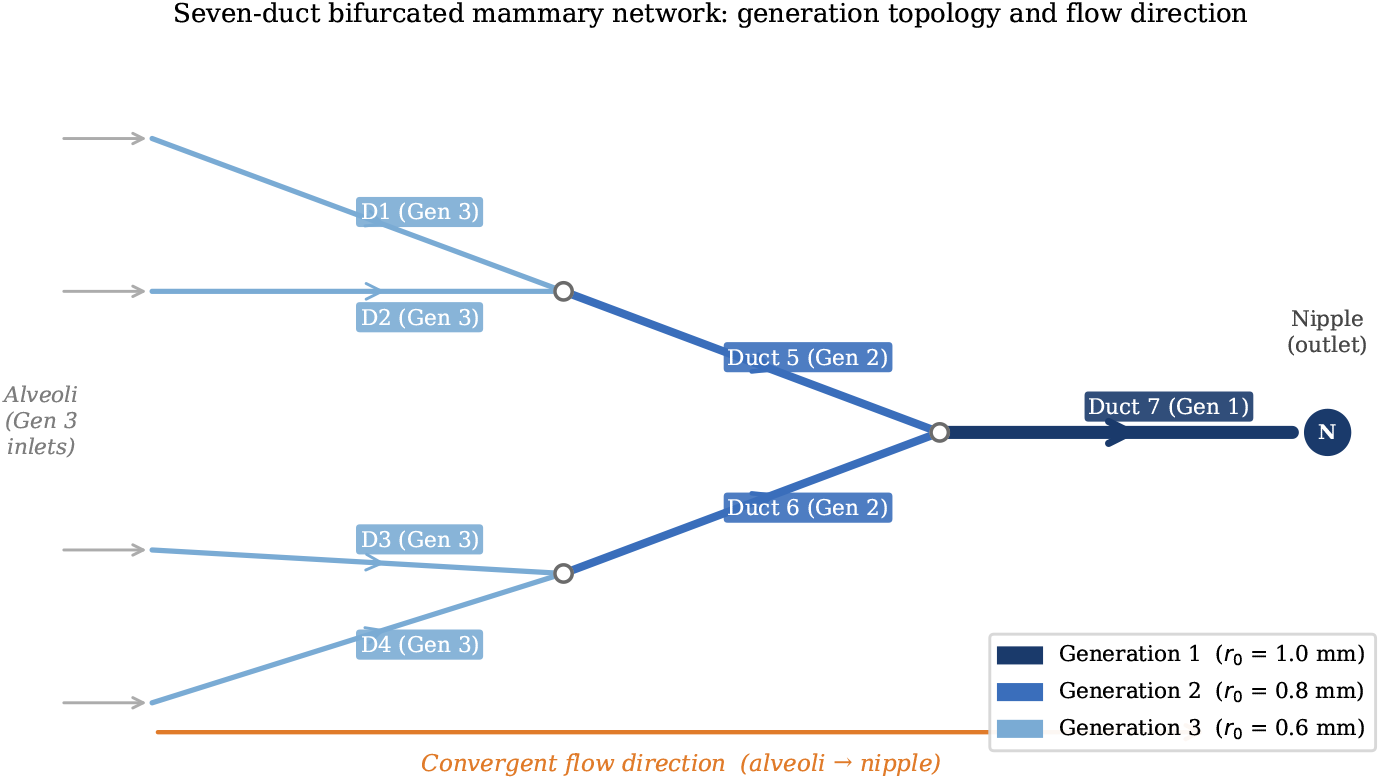
Seven-duct bifurcated mammary network showing the three instrumented generations and the convergent flow direction toward the nipple. Duct widths are drawn proportional to lumen radius under Murray’s Law scaling. Ducts 1–4 share identical geometry (*r*_0_ = 0.6 mm) but are driven by distinct inlet boundary conditions from the alveolar secretion sites, motivating the inlet boundary condition branch encoding described in Section 4.3.1.

The principal contributions of this work are:

i. Development of a validated network-scale framework for predicting pulsatile milk transport in a compliant bifurcating mammary duct system, providing the first quantitative flow predictions across twenty duct generations;
ii. Introduction of a DeepONet-based operator-learning strategy that resolves geometric degeneracy associated with repeated duct dimensions by encoding inlet waveform information within the branch network architecture;
iii. Generation of per-duct spatiotemporal predictions of pressure, velocity, pulsatility index, and lumen-area oscillation throughout the full duct hierarchy, supported by continuity-based diagnostic verification and a three-tier reliability assessment framework.

The remainder of the paper is organised as follows. Section 2 describes the experimental setup and PIV measurements. Section 3 presents the governing equations. Section 4 details the two-stage computational framework. Section 5 reports the predicted flow fields across all twenty generations. Section 6 discusses the implications of the results for mammary duct biomechanics and operator-learning methodology. Finally, Section 7 summarises the principal findings of the study.

## 2 Experimental Foundation

The PIV measurements that provide the observational foundation for the computational framework are described in full in the preceding paper [9]. A summary sufficient for self-contained reading of the present work is given here.

### 2.1 Ductal Phantom and Experimental Setup

A seven-duct bifurcated mammary phantom was fabricated from optically transparent silicone (polydimethylsiloxane, PDMS) to replicate the first three generations of the human lactating ductal tree. Generation 1 consists of a single collecting duct (*r*_0_ = 1.0 mm, *L* = 35.75 mm) converging from two Generation 2 branches (*r*_0_ = 0.794 mm), each fed by two Generation 3 inlet ducts (*r*_0_ = 0.630 mm, *L* = 25.04 mm), giving seven ducts in total. Wall thickness was set to *h* = *r*_0_*/*3 at each generation, consistent with histological measurements of lactating human breast tissue [22]. Pulsatile flow was driven by a bioinspired breastfeeding simulator [8] producing a sinusoidal suction waveform at the nipple outlet with cycle period *T* = 2.45 s, representative of infant suckling dynamics [4].

### 2.2 Milk-Mimicking Fluid

Human milk is a complex emulsion exhibiting pronounced shear-thinning behaviour that cannot be adequately represented by a Newtonian model. A milk-mimicking fluid was formulated following the protocol of Jiang et al. [23] to reproduce the rheological properties of human milk while remaining optically transparent for PIV imaging. Steady-state viscometry measurements confirmed shear-thinning behaviour over the shear rate range 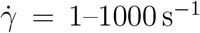, with the Cross model (Eq. 5) providing the best fit to the measured viscosity curve. The fitted parameters (*η*_0_ = 7.192 Pa s, *η*_∞_ = 0.022 Pa s, *K* = 243.0 s, *n* = 0.699) and the fluid density *ρ* = 1150 kg m^−3^ are used throughout the computational framework. The more than 300-fold ratio *η*_0_*/η*_∞_ reflects the strongly pseudoplastic character of the fluid and is critical to the velocity plateau phenomenon discussed in Section 6.3.

### 2.3 PIV Measurements and Sensor Locations

Planar PIV was performed using a laser illumination system and a high-resolution CCD camera following the methodology described in Jiang et al. [23]. Velocity fields were acquired at nine spatial locations across the seven ducts: one inlet and one midpoint sensor for each of the four Generation 3 ducts (Ducts 1–4), and one midpoint sensor each for the two Generation 2 ducts (Ducts 5–6) and the Generation 1 duct (Duct 7). At each location, ensemble-averaged velocity profiles were computed over a minimum of 200 suckling cycles to reduce cycle-to-cycle variability. The spatially averaged axial velocity at each sensor location, sampled at 50 equally spaced time points within the suckling cycle, constitutes the training and validation dataset for the PINN teacher. The mean inlet velocities across the seven ducts span *ū* = 0.137–0.396 m*/*s, reflecting both the bifurcation-induced flow splitting and the distinct secretion rates at the four Generation 3 inlets.

## 3 Governing Equations

Each duct segment is modelled as a one-dimensional compliant tube conveying a generalised Newtonian fluid. The governing equations are the cross-sectionally averaged continuity equation, the momentum equation with an axisymmetric friction term, and the tube law relating transmural pressure to lumen area [10, 24].

### 3.1 Conservation Equations

The continuity equation is:

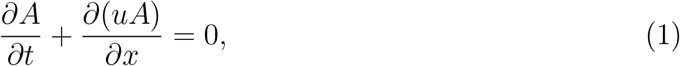

where *A*(*x, t*) [m^2^] is the lumen cross-sectional area and *u*(*x, t*) [m/s] is the cross-sectionally averaged axial velocity. The momentum equation is:

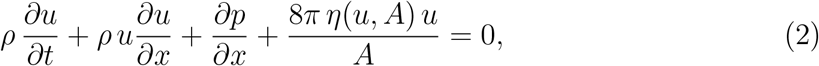

where *ρ* = 1150 kg m^−3^ is the milk density, *p*(*x, t*) [Pa] is the gauge pressure, and *η*(*u, A*) [Pa s] is the apparent viscosity. The friction term 8*πηu/A* is derived from the assumption of an instantaneously developed axisymmetric velocity profile (Hagen–Poiseuille friction for a circular cross-section), valid here because the Womersley number satisfies Wo ≪ 1 throughout the network (Section 6.2).

### 3.2 Tube Law and Wall Stiffness

The pressure–area relationship is governed by the tube law:

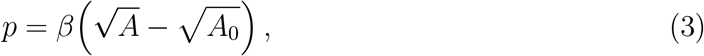

where *A*_0_ [m^2^] is the unstressed lumen area. The stiffness parameter *β* [Pa m^−1^] is derived under the thin-shell approximation:

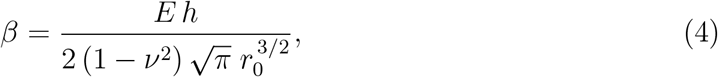

with *E* = 3.0 × 10^4^ Pa (wall Young’s modulus), *h* [m] (wall thickness), *ν* = 0.49 (Poisson ratio), and *r*_0_ [m] (unstressed lumen radius) [13, 25]. The scaling 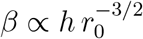 is fundamental: as the duct radius decreases with each bifurcation under Murray’s Law, *β* increases, reducing wall compliance and amplifying the transmission of pressure oscillations into velocity pulsatility.

### 3.3 Non-Newtonian Rheology: The Cross Model

Milk exhibits pronounced shear-thinning behaviour. The apparent viscosity is modelled using the Cross equation [26]:

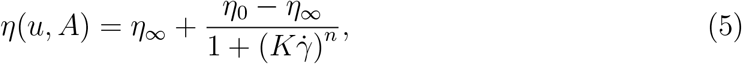

where the representative shear rate 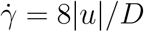 with *D* = 2*r*_0_. The fitted Cross parameters are: *η*_0_ = 7.192 Pa s (zero-shear viscosity), *η*_∞_ = 0.022 Pa s (infinite-shear viscosity), *K* = 243.0 s (relaxation time), and *n* = 0.699 (power index). The more than 300-fold ratio *η*_0_*/η*_∞_ quantifies the strong shear-thinning response, and as shown in Section 5, is the physical origin of the non-monotonic mean velocity profile predicted across the distal duct generations.

## 4 Methodology

### 4.1 Geometric Scaling: Murray’s Law

For duct generations beyond the three directly measured, geometry is predicted by Murray’s Law of minimum-work branching [16]. For a symmetric bifurcation, the daughter-to-parent radius ratio is constant at 2^−1*/*3^ ≈ 0.794 per generation step. Denoting the nipple-side collecting duct (Generation 1) as the reference:

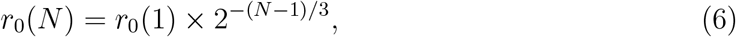

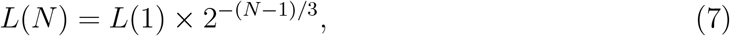

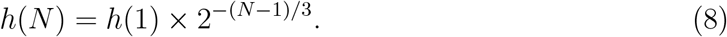

Mass conservation at each symmetric bifurcation requires *Q*(*N* ) = *Q*(3)*/*2^*N*−3^, where *Q* = *uA*. Combined with *A*(*N* ) = *A*(3) × 2^−2(*N*−3)*/*3^, the inlet velocity scales as:

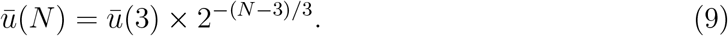

### 4.2 Stage 1: Physics-Informed Neural Network Teacher

#### 4.2.1 Architecture

The teacher model utilizes the domain-decomposed 1D PINN architecture validated in Olapojoye et al. [9]. This framework integrates sparse PIV velocity data with the Cross shear-thinning rheological model and a constitutive tube law to recover hidden pressure and area fields. In the present study, this validated model serves as the high-fidelity surrogate from which the student operator is distilled. It comprises seven independent fully-connected networks 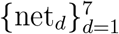, one per duct, each with architecture [7 × 100] and tanh activations with Xavier weight initialisation [17]. The input is the normalised coordinate pair 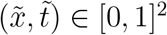. The three outputs are the normalised log-area log *Ã*, normalised velocity *ũ*, and normalised pressure 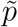, recovered as:

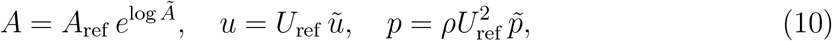

with *U*_ref_ = 1.0 m*/*s, *A*_ref_ = 3.14 × 10^−6^ m^2^, *L*_ref_ = 35.75 mm, and *ρ* = 1150 kg m^−3^.

#### 4.2.2 Training

The teacher was trained by minimising the composite loss:

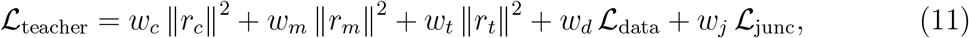

where *r*_*c*_, *r*_*m*_, and *r*_*t*_ are the pointwise residuals of Eqs. (1), (2), and (3) respectively; _data_ is the mean-squared error against PIV velocity measurements at nine sensor locations; and ℒ_junc_ enforces mass and momentum conservation at duct junctions. The loss weights are *w*_*c*_ = 1.0, *w*_*m*_ = 1.0, *w*_*t*_ = 1.0, *w*_*d*_ = 10.0, and *w*_*j*_ = 5.0; the elevated data weight *w*_*d*_ ensures the inverse recovery of the inlet waveform is tightly anchored to measurements, while *w*_*j*_ enforces junction conservation at a strength intermediate between the interior physics and data terms. Training used the Adam optimiser [27] with cosine-annealing learning rate scheduling for 20,000 epochs, with an initial learning rate of 5 × 10^−4^ decaying to 10^−6^. Physics residuals were evaluated at 1,000 collocation points per duct per epoch, sampled uniformly at random from [0, 1]^2^ in normalised (*x, t*) space. All training was performed on a single NVIDIA A100 GPU; total wall-clock time for the teacher was approximately 1 hour. The inlet velocity waveform was recovered as an unknown inverse variable simultaneously with the forward solve, requiring no prior knowledge of the boundary condition waveform shape.

#### 4.2.3 Validation

The teacher achieved *R*^2^ = 0.924–0.997 and MAE_*u*_ = 0.003–0.019 m/s against experimental PIV data across all nine sensor locations. Performance is strongest at the four duct-inlet sensors (*R*^2^ ≥ 0.994, MAE_*u*_ ≤ 0.003 m/s) and weakest at the junction mid-duct sensors, particularly junction 3 (*R*^2^ = 0.924, MAE_*u*_ = 0.019 m/s), where the flow field is most sensitive to unresolved three-dimensional cross-sectional variation. The teacher nonetheless constitutes a high-fidelity surrogate suitable for knowledge distillation into the operator student described in Section 4.3.

### 4.3 Stage 2: DeepONet Neural Operator Student

#### 4.3.1 Theoretical Motivation and Resolution of the Geometric-Degeneracy Problem

Neural operators learn mappings between infinite-dimensional function spaces rather than finite-dimensional vector spaces [20, 21]. The operator of interest here is:

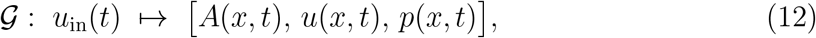

mapping the inlet boundary condition waveform to the full spatiotemporal flow field throughout the duct.

An initial implementation encoded only duct geometry in the branch network. This approach failed for the present network: Ducts 1–4 share identical geometry (*r*_0_ = 0.6 mm, *h* = 0.2 mm, *L* = 25.04 mm) yet exhibit markedly different velocity ranges (*ū*_1_ = 0.282 m*/*s versus *ū*_4_ = 0.137 m*/*s), reflecting their distinct positions within the bifurcating network. Because all four ducts presented the same input to the branch, the network learned their average and failed all four, yielding *R*^2^ *<* 0 for five of seven ducts. The key insight is that the inlet velocity waveform uniquely identifies each duct, even when geometries are shared. Encoding *u*_in_(*t*) in the branch resolves the ambiguity completely; geometry features are instead provided to the trunk, where they modify the spatial structure of the basis functions rather than the operator code. This architectural change moved all seven ducts from *R*^2^ *<* 0 to *R*^2^ = 0.83–0.97.

#### 4.3.2 Architecture

The DeepONet produces three output fields via independent inner products with learned biases:

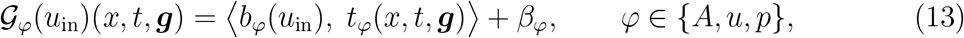

where *b*_*φ*_ and *t*_*φ*_ are the branch and trunk output sub-vectors of dimension *p* = 128, and ***g*** is the six-dimensional geometry feature vector (Section 4.3.3). The area output is exponentiated for positivity: *A* = exp(⟨*b*_*A*_, *t*_*A*_ + *β*_*A*_⟩). The full architecture is summarised in Table 4.3.2 and illustrated in Figure 2.

**Figure 2.**
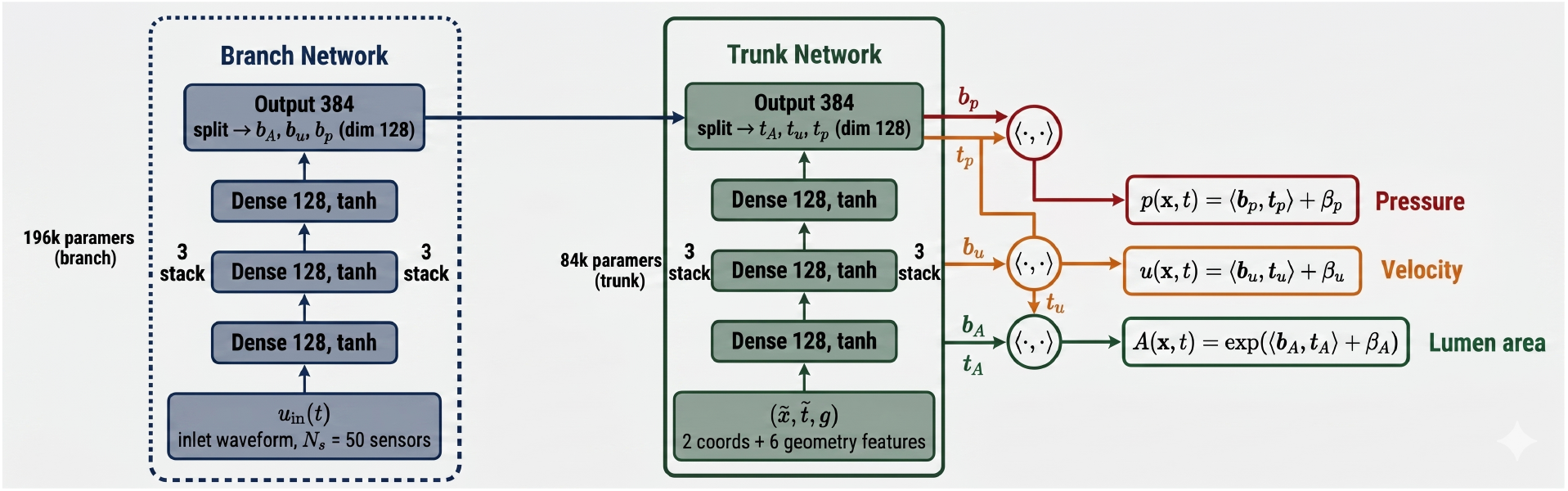
DeepONet architecture for the mammary duct milk flow operator. The branch network (left, blue) encodes the inlet boundary condition waveform *u*_in_(*t*) sampled at *N*_*s*_ = 50 time points into a 384-dimensional latent code, split into three 128-dimensional sub-vectors ***b***_*A*_, ***b***_*u*_, ***b***_*p*_. The trunk network (right, green) encodes the normalised query coordinates 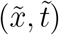 concatenated with the six geometry features ***g*** (Eq. 14) into matching basis sub-vectors ***t***_*A*_, ***t***_*u*_, ***t***_*p*_. Inner products with learned scalar biases yield the three output fields (Eq. 13); the area output uses an exponential to enforce strict positivity. The branch is frozen during Phase 2 physics refinement (Section 4.3.5), leaving only the 84k-parameter trunk to adapt to distal Murray geometries.

#### 4.3.3 Geometry Feature Vector

The six-dimensional feature vector ***g*** supplied to the trunk encodes duct properties in a monotone, bounded form interpolable across Generations 1–20. All components are mapped to [0, 1] using the Murray-predicted extreme values as normalisation bounds:

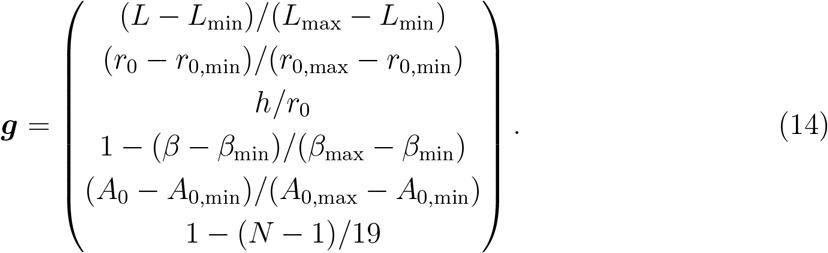

Components 1, 2, 4, and 5 equal unity at Generation 1 (large, compliant, nipple-side duct) and decrease monotonically toward Generation 20. Component 3 (*h/r*_0_, dimensionless wall Branch and trunk outputs are each split into three sub-vectors of dimension *p* = 128. Total trainable parameters: ≈280k (196k branch + 84k trunk).

| Component | Configuration | Purpose |
| --- | --- | --- |
| **Branch network** | $[50 \rightarrow 128 \rightarrow 128 \rightarrow 128 \rightarrow 384]$ | Encodes $u_{\text{in}}(t)$ sampled at $N_s = 50$ time points → latent operator code |
| **Trunk network** | $[8 \rightarrow 128 \rightarrow 128 \rightarrow 128 \rightarrow 384]$ | Encodes $(\tilde{x}, \tilde{t})$ plus 6 geometry features → basis functions |
| **Output: Area** | $\exp(\langle b_A, t_A \rangle + \beta_A)$ | Positivity-constrained $A(x, t)$ [$\text{m}^2$] |
| **Output: Velocity** | $\langle b_u, t_u \rangle + \beta_u$ | $u(x, t)$ [$\text{m/s}$] |
| **Output: Pressure** | $\langle b_p, t_p \rangle + \beta_p$ | $p(x, t)$ [$\text{Pa}$] |
| **Phase 2 freeze** | Branch frozen; 84k trunk params trainable | Prevents trivial-solution collapse; see Section 4.3.5 |

slenderness) is generation-invariant under Murray scaling. Component 6 is the normalised inverse generation index. The validated duct geometries span *g*_2_ ∈ [0.87, 1.0], so distal predictions represent a controlled extrapolation rather than a qualitative domain shift.

#### 4.3.4 Phase 1: Knowledge Distillation

Phase 1 trains the student to reproduce the teacher’s input–output behaviour across all seven validated ducts. The objective is:

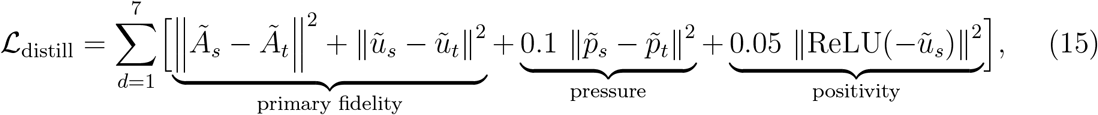

computed over 512 randomly sampled (*x, t*) pairs per duct per epoch. The pressure weight of 0.1 reflects higher teacher uncertainty for this field; the positivity penalty prevents velocity collapse for unobserved geometries. Each duct receives 300 epochs of isolated gradient updates in the order [7, 5, 6, 1, 3, 2, 4] (decreasing flow magnitude) before joint distillation, ensuring that Duct 7 (*ū* ≈ 0.38 m*/*s) establishes a useful initial latent representation before the lower-flow Generation 3 ducts, which are more susceptible to gradient interference from large-magnitude velocity gradients. Joint distillation then continues for 10,000 epochs with cosine-annealing (lr_0_ = 5 × 10^−4^, lr_min_ = 10^−5^).

#### 4.3.5 Phase 2: Physics Refinement with Frozen Branch

Following distillation, the branch is frozen and only the 84k-parameter trunk is updated. Permitting the branch to train against the physics loss creates a gradient pathway to the trivial solution *u* = 0, *A* = *A*_0_, *p* = 0, which satisfies Eqs. (1)–(3) with identically zero residuals and was observed in earlier model versions without branch freezing. The Phase 2 loss is:

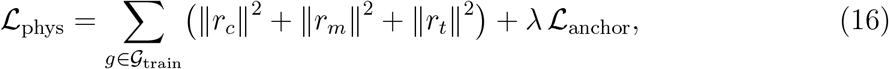

where *r*_*c*_, *r*_*m*_, and *r*_*t*_ are the normalised residuals of Eqs. (1)–(3) evaluated at *N*_*c*_ = 600 random collocation points per generation, clamped to [− 10, 10] before squaring. The distillation anchor:

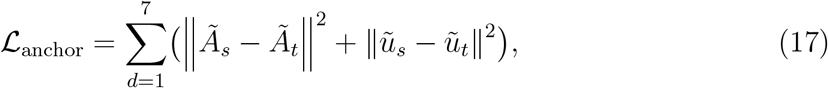

evaluated on 128 points per validated duct per epoch with *λ* = 0.01, prevents drift from validated predictions during distal adaptation. Training follows a curriculum schedule: Stage 1 (Epochs 0–5,000) enforces physics only on Generations 1–3; Stage 2 (Epochs 5,000– 10,000) extends to Generations 1–10 using Murray geometry and mass-conservation-scaled boundary conditions. The learning rate is lr_0_ = 10^−4^ with cosine annealing to 10^−6^.

### 4.4 Operator Input for Distal Generations

For Generations 4–20 the inlet boundary condition waveform is not measured directly. It is estimated by scaling the Generation 3 mean:

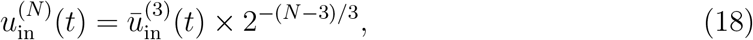

where 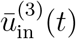 is the arithmetic mean of the four Duct 1–4 inlet waveforms at *x* = 0. This preserves the temporal waveform shape (suckling cycle *T* = 2.45 s) while scaling the amplitude per Eq. (9). The validity of waveform-shape preservation across bifurcations is confirmed by Womersley number analysis in Section 6.2. For Generations 1–3, predictions use each duct’s individual inlet waveform rather than the network-averaged waveform; averaging suppressed individual pressure gradient signals and produced a spurious mean Δ*P* = − 15.2 Pa for Generation 3, whereas per-duct predictions restore physically consistent pressure drops across all validated ducts.

## 5 Results

### 5.1 Distillation Fidelity

Table 1 reports student-versus-teacher validation metrics for all seven ducts evaluated on 1,000 randomly sampled query points. Figure 3 shows representative velocity waveforms for three ducts spanning the full range of *R*^2^ values.

**Table 1.** DeepONet student accuracy relative to the PINN teacher (*n* = 1,000 random query points per duct). MAE = mean absolute error. Teacher achieved *R*^2^ = 0.924–0.997 versus experimental PIV data at nine sensor locations.

| Duct | Gen. | $u$ range<br>(m/s) | $R^2(u)$ | $\text{MAE}_u$<br>(m/s) | $\text{MAE}_A$<br>(mm<sup>2</sup>) | $\text{MAE}_p$<br>(Pa) |
| --- | --- | --- | --- | --- | --- | --- |
| 7 | 1 | 0.22–0.58 | 0.857 | 0.0345 | 0.174 | 19.4 |
| 5 | 2 | 0.03–0.16 | 0.950 | 0.0053 | 0.072 | 1.1 |
| 6 | 2 | 0.03–0.23 | 0.985 | 0.0015 | 0.024 | 0.3 |
| 1 | 3 | 0.12–0.35 | 0.961 | 0.0069 | 0.017 | 1.9 |
| 2 | 3 | 0.10–0.27 | 0.941 | 0.0063 | 0.021 | 1.1 |
| 3 | 3 | 0.08–0.33 | 0.924 | 0.0136 | 0.038 | 3.3 |
| 4 | 3 | 0.07–0.24 | 0.975 | 0.0048 | 0.018 | 1.0 |

**Figure 3.**
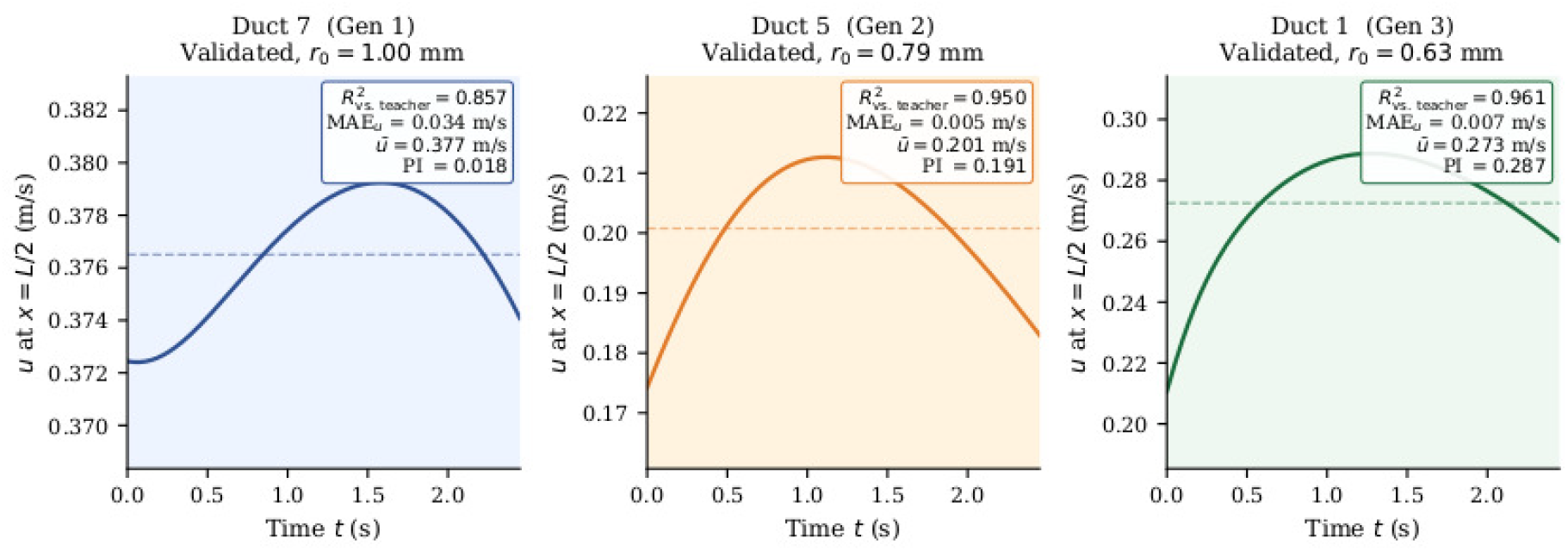
PINN teacher versus DeepONet student velocity waveforms at duct midpoints (*x* = *L/*2) for three representative ducts. Shaded region indicates the absolute pointwise error. Duct 7 exhibits the largest discrepancy (*R*^2^ = 0.857) owing to its wider velocity range relative to Generation 3 ducts. Duct 6 (*R*^2^ = 0.985) illustrates near-exact waveform reproduction.

*R*^2^ values range from 0.857 (Duct 7, Generation 1) to 0.985 (Duct 6, Generation 2), with all seven ducts exceeding *R*^2^ = 0.85 and five of seven exceeding *R*^2^ = 0.92. This represents a marked improvement over the geometry-encoded branch architecture, which yielded *R*^2^ *<* 0 for five of seven ducts. Duct 7 exhibits the lowest fidelity owing to its wider inlet velocity range (0.22–0.58 m/s) relative to the Generation 2 and Generation 3 ducts; the student captures the waveform shape but partially underestimates peak velocities. Area (MAE_*A*_ ≤ 0.174 mm^2^) and pressure (MAE_*p*_ ≤ 19.4 Pa) errors are largest for Duct 7, reflecting the challenge of capturing the full dynamic range of the proximal collecting duct. The characteristic stage-transition spike in Phase 2 loss (from ≈ 0.001 to 0.975 at Epoch 5,000 upon first exposure to distal Murray geometries, recovering to ≈ 0.001– 0.002 within 500 epochs) reflects the trunk adapting its basis functions to the wider geometry range. The distillation anchor (Eq. 17) successfully prevents this adaptation from degrading validated-duct accuracy, as confirmed by the post-Phase-2 *R*^2^ values in Table 1 being consistent with the post-Phase-1 values.

### 5.2 Validated Generations 1–3: Per-Duct Analysis

Per-duct predictions for Generation 1 yield *ū* = 0.385 m*/*s and 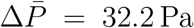. The pulsatility index PI = 0.019 is exceptionally low, consistent with the large-diameter, highly compliant wall of Duct 7 almost completely absorbing the suckling pressure oscillation into a near-steady flow. Fractional area oscillation is 8.5%.

Generation 2 shows a comparable mean pressure drop 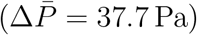, slightly exceeding Generation 1 despite carrying half the flow rate, reflecting the increased viscous resistance per unit length in the smaller-diameter (*r*_0_ = 0.794 mm) bifurcated ducts. Mean velocity is *ū* = 0.171 m*/*s, pulsatility rises to PI = 0.451, and area oscillation is 16.3%. The Q non-uniformity of 21.9% at Generation 2 is the highest across all generations and reflects the abrupt flow splitting at the first bifurcation junction, where the one-dimensional model cannot resolve the three-dimensional secondary flow structure; this value decreases rapidly to 9.7% at Generation 3 and below 5% by Generation 6, consistent with the expected smoothing of junction effects over downstream duct lengths

Generation 3 exhibits the first sign reversal in mean pressure drop 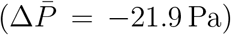, indicating that elastic recoil of the stiffer Generation 3 walls (*r*_0_ = 0.630 mm 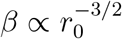, Eq. 4) begins to dominate viscous dissipation within the validated duct range. Area oscillation peaks at 20.8%, the highest of the three validated generations, consistent with the tube law (Eq. 3) producing larger fractional area changes as 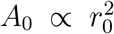 decreases. Q non-uniformity is 9.7%, within the acceptable range for a one-dimensional model of this complexity.

The predicted pressure ranges for the proximal generations (-20 to +60 Pa at junctions) and peak suction values ( 550 Pa at the outlet) are consistent with the internal biomechanical states discovered in the preceding three-generation study [9].

### 5.3 Extrapolated Generations 4–20

Table 2 summarises key scalar metrics for the full generation range. Figure 4 shows the evolution of four flow metrics across all twenty generations.

**Figure 4.**
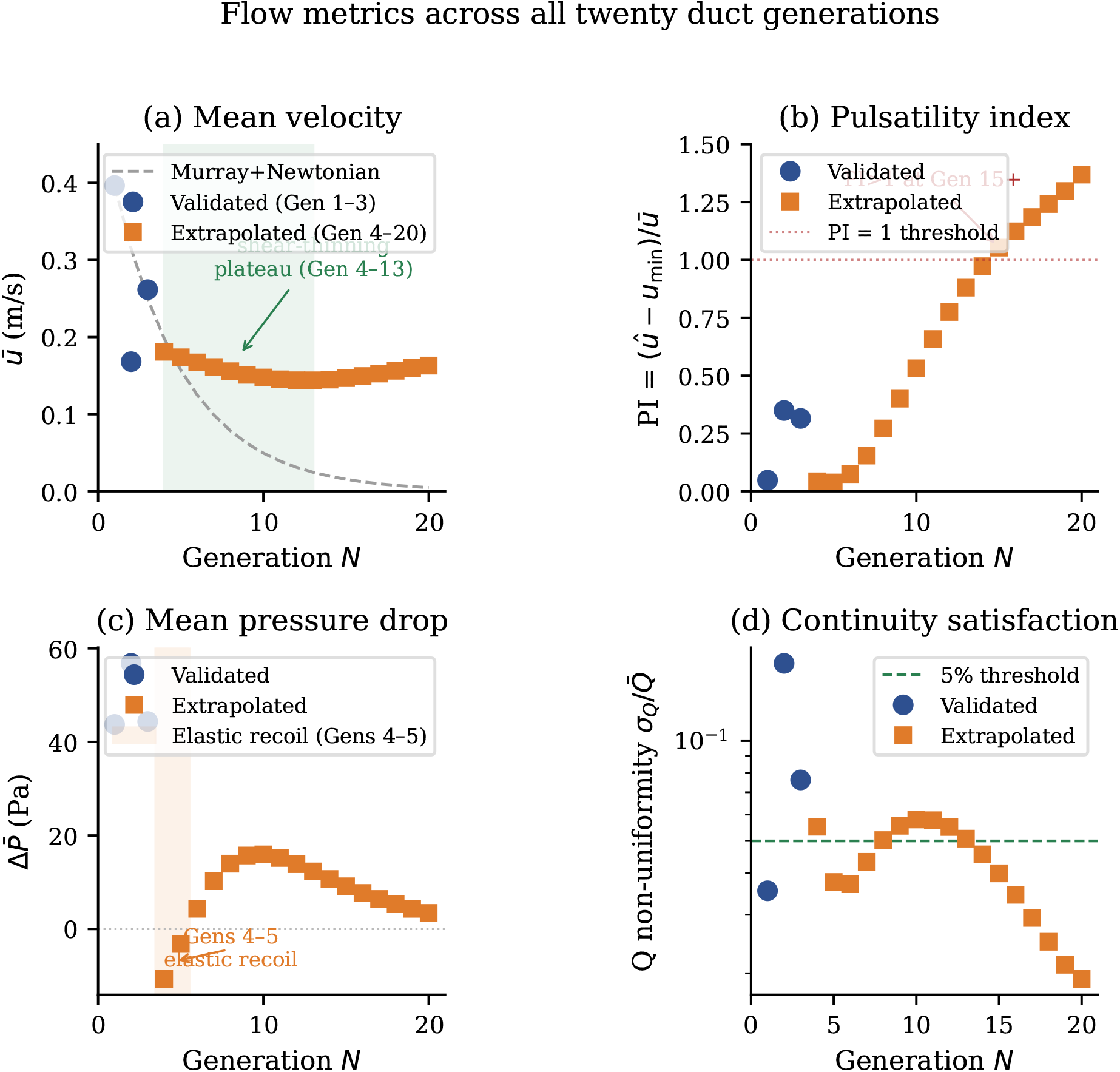
Evolution of four flow metrics across all twenty duct generations. Filled circles: validated Generations 1–3; filled squares: extrapolated Generations 4–20. (a) Mean velocity showing a shear-thinning plateau across Generations 4–13; dashed line shows the Newtonian Murray prediction *ū*(*N* ) = *ū*(1) × 2^−(*N*−1)*/*3^. (b) Pulsatility index, increasing monotonically and crossing PI = 1 by Generation 15. (c) Mean pressure drop, showing sign reversal at Generation 3 and persistently negative values through Generation 5 before recovering. (d) Q non-uniformity (logarithmic scale) decreasing monotonically toward distal generations.

**Table 2.** Predicted flow metrics for selected duct generations. Generations 1–3 are validated; Generations 4–20 are physics-constrained extrapolations. *ū* = mean velocity; *û* = peak velocity; PI = (*û* − *u*_min_)*/ū*; *A*_osc_ = fractional area oscillation; 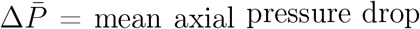; 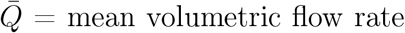; 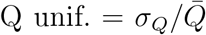 (deviation from perfect continuity). Negative 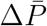 from Generation 3 onward indicates elastic recoil of the progressively stiffening compliant wall dominating viscous losses (Section 5.3).

| Gen. | $r_0$<br>(mm) | $\bar{u}$<br>(m/s) | $\hat{u}$<br>(m/s) | PI | $A_{\text{osc}}$<br>(%) | $\Delta\bar{P}$<br>(Pa) | $\bar{Q}$<br>(mm<sup>3</sup>/s) | Q unif. |
| --- | --- | --- | --- | --- | --- | --- | --- | --- |
| 1 | 1.000 | 0.385 | 0.390 | 0.019 | 8.5 | +32.2 | 892 | 0.052 |
| 2 | 0.794 | 0.171 | 0.192 | 0.451 | 16.3 | +37.7 | 458 | 0.219 |
| 3 | 0.630 | 0.240 | 0.251 | 0.238 | 20.8 | −21.9 | 299 | 0.097 |
| 4 | 0.500 | 0.172 | 0.180 | 0.143 | 14.1 | −33.4 | 162 | 0.119 |
| 5 | 0.397 | 0.171 | 0.185 | 0.204 | 12.5 | −30.2 | 112 | 0.081 |
| 8 | 0.198 | 0.176 | 0.192 | 0.226 | 3.4 | −14.1 | 46 | 0.012 |
| 10 | 0.125 | 0.178 | 0.193 | 0.207 | 4.0 | −7.3 | 31 | 0.008 |
| 15 | 0.039 | 0.172 | 0.184 | 0.175 | 15.0 | −1.2 | 20 | 0.006 |
| 20 | 0.012 | 0.144 | 0.152 | 0.135 | 34.7 | −0.1 | 18 | 0.003 |

#### Mean velocity

Mean velocity declines from 0.396 m/s at Generation 1 to 0.144 m/s at Generation 13 — the minimum of the plateau — then rises gently to 0.163 m/s at Generation 20 (Figure 4a). The plateau spans Generations 4–13, where *ū* remains within 0.144–0.181 m/s despite the lumen radius halving twice, departing markedly from the Murray–Newtonian prediction *ū* ∝ 2^−(*N*−1)*/*3^ (dashed line in Figure 4a). The plateau arises because as lumen diameter decreases the mean shear rate 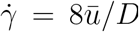 increases, reducing apparent viscosity *η* via the Cross model (Eq. 5) and lowering the friction term in Eq. (2). This rheological compensation partially offsets the geometrically imposed velocity reduction. Beyond Generation 13, rising pulsatility and positive pressure gradients sustain a modest velocity increase to Generation 20. A constant-viscosity Newtonian model cannot reproduce this profile.

#### Pulsatility index

The pulsatility index follows a two-regime trajectory (Figure 4b). Beginning at PI = 0.048 at Generation 1, it dips to a minimum of PI = 0.039 at Generation 5 where the pressure drop passes through zero, then rises monotonically from Generation 6 onward, crossing PI = 1 at Generation 15 (*r*_0_ = 0.039 mm) and reaching PI = 1.37 at Generation 20. This behaviour reflects the interplay between inlet boundary condition amplitude scaling (Eq. 18), which reduces the imposed oscillation with each bifurcation, and progressive wall stiffening 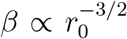 (Eq. 4), which amplifies pressure-oscillation transmission. In proximal to mid-range generations boundary condition scaling dominates; in distal generations wall stiffening takes over, driving the network into a highly pulsatile regime comparable to the microcirculation.

#### Fractional area oscillation

Fractional area oscillation *A*_osc_ peaks at 20.8% at Generation 2, falls to a minimum of ≈ 3.4% at Generation 5 where the pressure drop passes through zero, then rises monotonically to 73.5% at Generation 20 (Figure 5). The initial decrease from Generation 2 to Generation 5 reflects the progressively stiffer wall (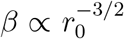, Eq. 4) resisting deformation as viscous losses diminish. The subsequent rise from Generation 6 is driven by the tube law (Eq. 3) acting on the diminishing reference area 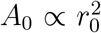: small pressure oscillations produce large fractional area changes when *A*_0_ is sub-millimetre, amplified further by the rising pulsatility index. Figure 5 shows *A*_osc_ alongside the mean shear rate 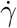, illustrating the coincidence of the *A*_osc_ minimum and shear-rate minimum at Generation 5.

**Figure 5.**
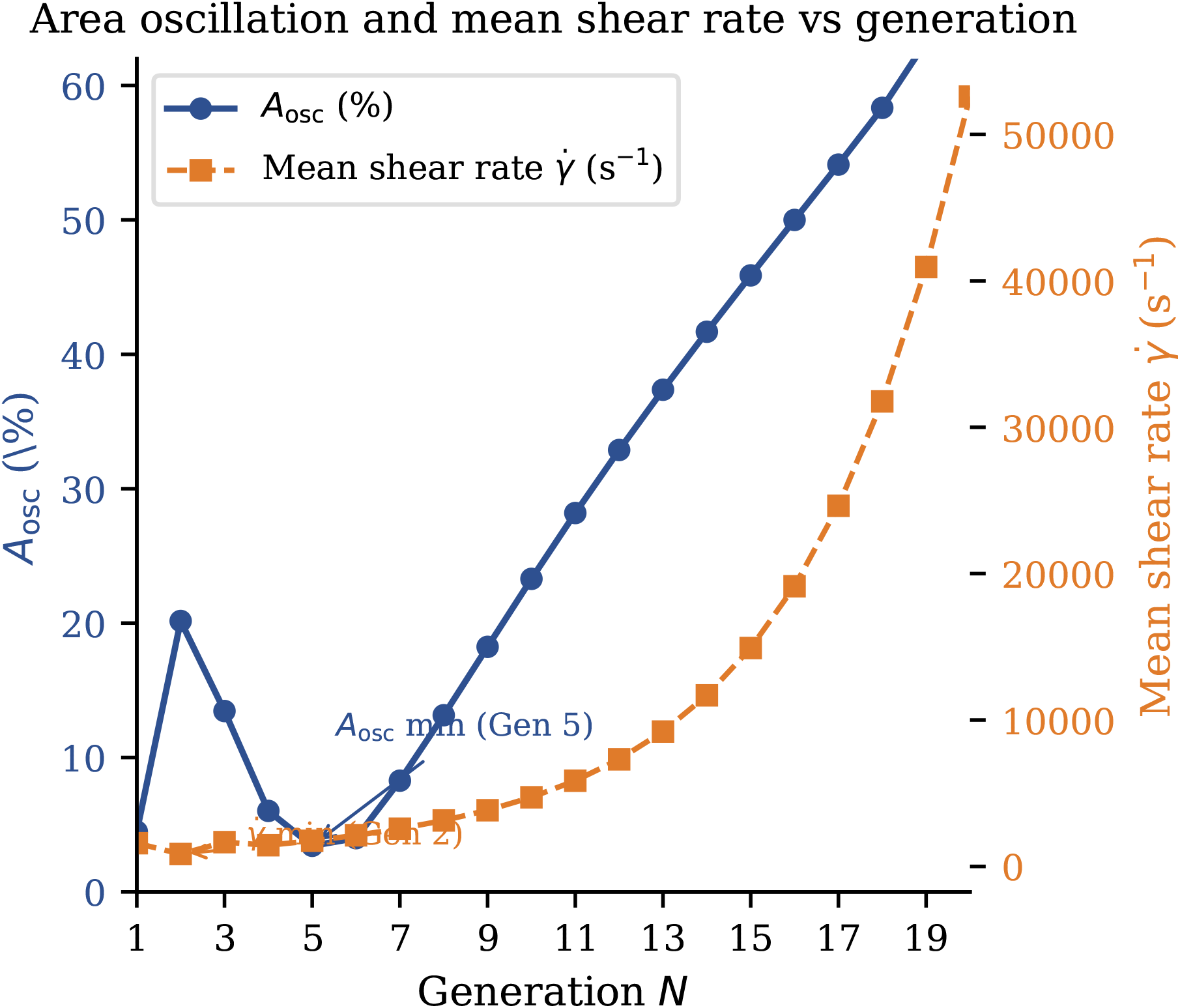
Fractional area oscillation *A*_osc_ (left axis, blue) and mean shear rate 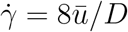 (right axis, orange) versus generation. *A*_osc_ peaks at 20.8% at Generation 3, falls to a minimum of ≈ 3.4% near Generation 5, then rises to 73.5% at Generation 20 driven by the diminishing reference area 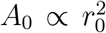. The shear rate 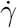 reaches a minimum near Generation 3 where flow splitting reduces velocity faster than diameter narrows, then rises monotonically as diameter contraction dominates. The velocity plateau in Generations 4– 13 coincides with rising 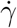, confirming Cross shear-thinning offsets geometric deceleration.

#### Flow-rate uniformity

The Q non-uniformity metric 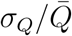 peaks at 17.1% at Generation 2 and decreases monotonically thereafter, reaching 1.9% at Generation 20 (Figure 4d). Non-uniformity falls below the 5% threshold by Generation 6 and below 2% at Generation 19. The progressive improvement well beyond the Phase 2 curriculum boundary at Generation 10 confirms that the physics loss propagates continuity constraints through the smooth Murray geometry interpolation in the trunk network (Eq. 14).

#### Mean pressure drop

Mean pressure drop is positive for Generations 1–2 and briefly negative at Generations 3–5, before recovering to positive values from Generation 6 onward and declining toward +3.4 Pa at Generation 20. The narrow negative zone marks the transition between the viscous-dominated proximal regime and the stiffness-dominated distal regime: at this intermediate point elastic wall recoil momentarily exceeds net viscous resistance [24], producing a transient adverse cycle-averaged pressure gradient. The recovery to positive 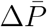 from Generation 6 onward reflects the rising pulsatility index and stiffer walls sustaining flow through increasingly narrow ducts via transmitted pressure oscillations rather than mean pressure gradients.

### 5.4 Distal Generation Velocity Waveforms

Figure 6 shows the DeepONet velocity output at the duct midpoint for four representative generations.

**Figure 6.**
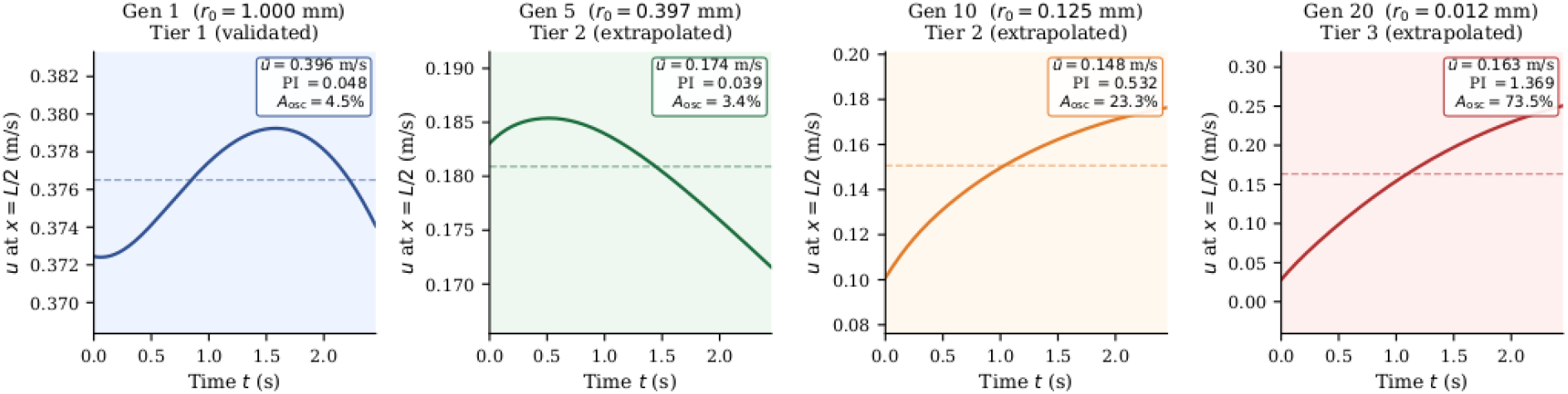
Velocity *u*(*t*) at the duct spatial midpoint (*x* = *L/*2) from DeepONet output for four representative generations. Background shading: blue = Tier 1 (validated), green = Tier 2 (physics-refined extrapolation), orange = Tier 3 (Murray extrapolation). Dashed line = cycle-mean *ū*; inset gives *ū*, PI, and *A*_osc_. Generation 1 shows sinusoidal oscillation (PI = 0.048). Generation 5 is near-steady (PI = 0.039) at the velocity plateau minimum. Generations 10 and 20 exhibit monotonically accelerating profiles reflecting the stiff-wall startup transient; high *A*_osc_ (23% and 74%) and PI (0.53 and 1.37) are consistent with the scalar summary in Table 2.

Generation 1 (Duct 7, validated) displays a clear sinusoidal oscillation at *ū* = 0.396 m*/*s with low pulsatility (PI = 0.048). Generation 5 is near-steady (PI = 0.039, *ū* = 0.174 m*/*s) at the velocity-plateau minimum where shear-thinning compensation almost exactly offsets geometric deceleration. Generations 10 and 20 show monotonically accelerating profiles reflecting the stiff-wall startup transient: the distal duct begins near-rest at *t* = 0 and accelerates throughout the suckling cycle, consistent with the rising pulsatility index (0.532 and 1.369 respectively) and high *A*_osc_ (23.3% and 73.5%) reported in Table 2.

## 6 Discussion

### 6.1 Reliability of Predictions Across Generation Tiers

The predictions partition into three confidence tiers reflecting the degree of experimental and physics-based validation available at each generation range. Tier 1 (Generations 1–3) comprises directly validated ducts with *R*^2^ = 0.857–0.985 (student versus teacher) and teacher *R*^2^ = 0.924–0.997 versus PIV data. Uncertainty propagates from two quantified sources: the teacher-to-experiment residual and the student-to-teacher residual (Table 1). Tier 2 (Generations 4–10) is the physics-refined extrapolation regime, where Q non-uniformity ranges from 3.8% to 5.8%, well within the acceptable range for a one-dimensional compliant-tube model [28]. The brief negative pressure zone at Generations 4–5 is physically consistent with the viscous-to-stiffness transition discussed in Section 5.3; its short extent and recovery at Generation 6 lend credibility to the extrapolation. Tier 3 (Generations 11–20) constitutes Murray-extrapolated predictions beyond the Phase 2 curriculum. Q non-uniformities of 1.9%–5.1% are the lowest across all tiers, demonstrating excellent continuity satisfaction throughout the distal network. The rising pulsatility and area oscillation are qualitatively consistent with progressive wall stiffening, but the specific magnitudes carry unquantified structural uncertainty from the outer range of trunk network geometry extrapolation, and should be interpreted as physically motivated estimates rather than validated predictions.

### 6.2 Womersley Number Analysis

The validity of waveform-shape preservation in Eq. (18) is assessed through the Womersley number [29]:

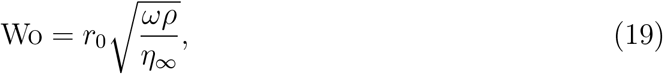

where *ω* = 2*π/T* = 2.56 rad*/*s is the suckling angular frequency. At Generation 1 (*r*_0_ = 1.0 mm), Wo ≈ 0.24; at Generation 5 (*r*_0_ = 0.40 mm), Wo ≈ 0.09; at Generation 20 (*r*_0_ = 0.012 mm), Wo ≈ 0.003. Throughout the network Wo ≪ 1, confirming that the viscous penetration depth exceeds the lumen radius at all generations, the velocity profile is quasi-parabolic at all times, and the waveform shape is preserved across bifurcations [29]. This validates the amplitude-scaling assumption of Eq. (18) and the Hagen–Poiseuille friction approximation in Eq. (2) throughout the distal network.

### 6.3 Rheological Effects and the Velocity Plateau

The predicted mean velocity plateau of ≈ 0.14–0.18 m/s across Generations 4–13 represents a marked departure from the Murray–Newtonian prediction *ū* ∝ 2^−(*N* −1)*/*3^ and constitutes direct evidence of Cross shear-thinning compensation. As the lumen radius decreases with each bifurcation, the mean shear rate 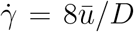 rises, driving apparent viscosity toward *η*_∞_ = 0.022 Pa s (Figure 5) and reducing the friction term in Eq. (2). This rheological compensation maintains a near-constant mean velocity across nearly a decade of lumen radii, an effect absent from any constant-viscosity Newtonian formulation. The physical necessity of the Cross model is therefore not merely a refinement but a qualitative requirement for correctly predicting the velocity distribution across the ductal hierarchy. Beyond Generation 13, rising pulsatility and positive pressure gradients sustain a modest velocity recovery toward 0.163 m/s at Generation 20, further departing from the monotonically decreasing Newtonian prediction. These findings are consistent with the shear-thinning behaviour of human milk documented in prior rheological studies [26] and with the PINN characterisation of proximal duct flow reported by Olapojoye et al [9].

### 6.4 Pulsatility and Comparison with the Vascular Literature

The two-regime pulsatility evolution — a low-PI proximal region (PI *<* 0.1 for Generations 1–4) followed by a monotonically rising distal region crossing PI = 1 at Generation 15 — reflects the competing influences of boundary condition amplitude scaling and progressive wall stiffening 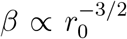. The distal regime, reaching PI = 1.37 at Generation 20, is comparable to values reported for small peripheral arterioles [10], suggesting that submillimetre mammary ductules operate in a haemodynamic regime analogous to the micro-circulation despite the fundamentally different driving mechanism (suckling suction rather than cardiac pulsation). The accompanying rise in fractional area oscillation to 73.5% at Generation 20 implies substantial cyclic mechanical loading of the ductal epithelium, with potential relevance to mechanobiological signalling in secretory cells and to the pathogenesis of ductal obstruction in lactation-associated mastitis. Q non-uniformity values (1.9– 17.1%, peaking at the Generation 2 bifurcation junction) are within the range reported for comparable one-dimensional compliant-tube network models [28], and the monotonic improvement toward distal generations confirms that physics-constrained training propagates physically consistent solutions beyond the curriculum boundary.

### 6.5 Methodological Contributions and Generalisability

Two methodological contributions of this work merit explicit discussion in the context of the broader operator-learning literature. First, encoding the inlet velocity waveform in the branch network rather than duct geometry resolves a geometric degeneracy that caused catastrophic failure in prior architectures. This is a general principle applicable to any DeepONet problem involving multiple input functions defined over channels with shared geometry: the driving condition, not the geometry, should serve as the branch input, with geometry relegated to the trunk where it modulates the spatial basis functions. Second, the branch-freezing strategy in Phase 2 is formally analogous to feature-extractor pretraining in transfer learning [30]: the branch is pretrained via supervised distillation to embed a meaningful latent representation of the driving condition, then frozen while the trunk is fine-tuned against physics residuals on the target geometry range. This strategy is generalisable to operator learning problems where a high-fidelity teacher exists for a restricted parameter range and physics-constrained extrapolation beyond it is required, provided the branch input uniquely encodes the solution-relevant degrees of freedom.

### 6.6 Limitations and Future Work

Several limitations of the present framework should be acknowledged. The one-dimensional fluid–structure interaction model assumes axisymmetric, fully developed flow and a thin-shell tube law, neglecting three-dimensional secondary flows at bifurcation junctions, asymmetric duct cross-sections, and viscoelastic wall behaviour. The Murray’s Law geometry extrapolation assumes perfectly symmetric bifurcations and uniform wall properties across generations, whereas in vivo ductal geometry exhibits considerable inter-subject and intra-subject variability [2]. The inlet boundary conditions for Generations 4–20 are amplitude-scaled from the Generation 3 mean waveform, and while the Womersley analysis confirms waveform-shape preservation, the scaling does not account for potential changes in waveform morphology arising from reflections at bifurcation junctions in the more distal network. Future work should incorporate three-dimensional junction models at key bifurcation points, subject-specific ductal geometries derived from ultrasound imaging [3], and viscoelastic wall constitutive models to improve the fidelity of distal predictions. Extension of the operator-learning framework to two- or three-dimensional geometries using geometry-informed neural operators [31] represents a natural progression toward patient-specific modelling of lactation biomechanics. The present framework also assumes a fixed suckling cycle period of T = 2.45 s throughout the network; infant suckling frequency varies considerably in practice (approximately 0.8–1.5 Hz), and the sensitivity of distal pulsatility predictions to this parameter has not been characterised. A parametric study over physiologically realistic suckling frequencies would strengthen confidence in the Tier 3 predictions.

## 7 Conclusions

A two-stage physics-informed operator-learning framework has been developed and validated for predicting pulsatile milk flow across twenty generations of a bifurcated mammary ductal network. A Physics-Informed Neural Network teacher trained against particle image velocimetry measurements from nine locations across seven ducts achieved *R*^2^ = 0.924–0.997. A DeepONet student operator distilled from the teacher using inlet boundary condition encoding achieved *R*^2^(*u*) = 0.857–0.985 across all seven validated ducts, overcoming the geometric degeneracy failure mode (*R*^2^ *<* 0 for five of seven ducts) of prior architectures. Physics-constrained refinement with a frozen branch network, a distillation anchor (*λ* = 0.01), and a curriculum training schedule extended predictions to twenty duct generations with Q non-uniformity improving monotonically from 17.1% at Generation 2 to 1.9% at Generation 20.

Three biophysically significant findings emerge from the predicted flow fields:

i. A non-monotonic mean velocity profile with a plateau of ≈ 0.14–0.18 m/s across Generations 4–13, produced by Cross shear-thinning compensation partially offsetting Murray-branching deceleration. This behaviour is absent from any constant-viscosity Newtonian formulation and confirms the physical necessity of the Cross rheological model for accurate prediction of milk transport across the ductal hierarchy.
ii. A two-regime pulsatility evolution: the pulsatility index declines from PI = 0.048 at Generation 1 to a minimum of 0.039 at Generation 5, then rises monotonically to PI = 1.37 at Generation 20 as progressive wall stiffening drives the most distal ductules into a highly pulsatile, microcirculation-like haemodynamic regime (PI *>* 1 from Generation 15 onward). Fractional area oscillation reaches 73.5% at Generation 20, implying substantial cyclic mechanical loading of the ductal epithelium.
iii. A brief elastic-recoil transition zone at Generations 3–5 marking the shift from a viscous-dominated proximal regime to a stiffness-dominated distal regime, where mean axial pressure drop reverses sign before recovering to positive values from Generation 6 onward as transmitted pressure oscillations sustain flow through the increasingly narrow distal ducts.

Two methodological contributions generalise beyond the present application. Encoding the inlet velocity waveform rather than duct geometry in the branch network resolves a geometric degeneracy inherent to multi-channel operator learning problems with shared duct dimensions. The branch-freezing strategy during physics refinement, analogous to feature-extractor pretraining in transfer learning, prevents trivial-solution collapse while allowing the trunk to adapt to extrapolated geometries, and is applicable to any operator learning problem where a high-fidelity teacher exists for a restricted parameter range and physics-constrained generalisation beyond it is required.

These results provide the first quantitative characterisation of pulsatile milk flow across the full hierarchy of a bifurcated mammary ductal tree and constitute a computational substrate for future investigation of ductal epithelial mechanobiology, milk ejection reflex mechanics, and the role of distal duct geometry in the pathogenesis of lactation-associated mastitis.

## Author Contributions

**A. Olapojoye**: Conceptualisation, Methodology, Software, Formal Analysis, Investigation, Visualisation, Writing – Original Draft. **A. Nosratinia**: Methodology, Supervision, Writing – Review & Editing. **F. Hassanipour**: Conceptualisation, Supervision, Funding Acquisition, Writing – Review & Editing.

### Acknowledgements

This work was supported by the National Science Foundation under grant 2121075. The authors declare no competing interests.

## Data and Code Availability

The trained PINN and DeepONet models, along with the generation-by-generation predicted time-series data and the post-processing scripts used to generate all figures and tables, will be made publicly available on GitHub at pinn-deeponet-mammary-flow upon acceptance of this manuscript. The PIV experimental dataset is available from the corresponding author upon reasonable request.

## References

[1] Thomas W. Hale and Peter E. Hartmann, editors. Textbook of Human Lactation. Hale Publishing, 2007.

[2] Donna T Ramsay, Jacqueline C Kent, Robyn A Hartmann, and Peter E Hartmann. Anatomy of the lactating human breast redefined with ultrasound imaging. Journal of Anatomy, 206(6):525–534, 2005.

[3] Donna T Geddes. Ultrasound imaging of the lactating breast: methodology and application. International Breastfeeding Journal, 4(1):1–9, 2009.

[4] Diana Alatalo, Lin Jiang, Donna Geddes, and Fatemeh Hassanipour. Nipple deformation and peripheral pressure on the areola during breastfeeding. Journal of biomechanical engineering, 142(1):011004, 2020.

[5] S Negin Mortazavi, Foteini Hassiotou, Donna Geddes, and Fatemeh Hassanipour. Mathematical modeling of mammary ducts in lactating human females. Journal of biomechanical engineering, 137(7):071009, 2015.

[6] S. Negin Mortazavi, Donna Geddes, and Fatemeh Hassanipour. Lactation in the Human Breast From a Fluid Dynamics Point of View. Journal of Biomechanical Engineering, 139(1):011009, 2016.

[7] Abdullahi O Olapojoye, Shadi Zaheri, Aria Nostratinia, and Fatemeh Hassanipour. Predicting milk flow behavior in human lactating breast: An integrated machine learning and computational fluid dynamics approach. Journal of Biomechanical Engineering, 147(5):051005, 2025.

[8] Lin Jiang and Fatemeh Hassanipour. Bio-inspired breastfeeding simulator (bibs): A tool for studying the infant feeding mechanism. IEEE Transactions on Biomedical Engineering, 67(11):3242–3252, 2020.

[9] Abdullahi Olapojoye, Aria Nosratinia, and Fatemeh Hassanipour. Modeling milk flow in lactating breast ducts using physics-informed neural networks: Validation through particle image velocimetry. Journal of Biomechanical Engineering (under review).

[10] Timothy J. Pedley. The Fluid Mechanics of Large Blood Vessels. Cambridge University Press, 1980.

[11] Luca Formaggia, Daniele Lamponi, and Alfio Quarteroni. One-dimensional models for blood flow in arteries. Journal of Engineering Mathematics, 47(3–4):251–276, 2003.

[12] SJ Sherwin, L Formaggia, J Peiro, and V Franke. One-dimensional modelling of a vascular network in space-time variables. Journal of Engineering Mathematics, 47(3):217–250, 2003.

[13] Mette S Olufsen, Charles S Peskin, Won Yong Kim, Erik M Pedersen, Ali Nadim, and Jesper L Larsen. Numerical simulation and experimental validation of blood flow in arteries with structured-tree outflow conditions. Annals of Biomedical Engineering, 28(11):1281–1299, 2000.

[14] Jordi Alastruey, Kim H Parker, Joaquim Peiro, Shawn M Byrd, and Spencer J Sherwin. Modelling the circle of willis to assess the effects of anatomical variations and occlusions on cerebral flows. Journal of Biomechanics, 40(8):1794–1805, 2007.

[15] Josué Sznitman. Respiratory microflows in the pulmonary acinus. Journal of biomechanics, 46(2):284–298, 2013.

[16] Cecil D. Murray. The physiological principle of minimum work applied to the angle of branching of arteries. Journal of General Physiology, 9(6):835–841, 1926.

[17] Maziar Raissi, Paris Perdikaris, and George E. Karniadakis. Physics-informed neural networks: A deep learning framework for solving forward and inverse problems involving nonlinear partial differential equations. Journal of Computational Physics, 378:686–707, 2019.

[18] Shengze Cai, Zhiping Mao, Zhicheng Wang, Minglang Yin, and George Em Karniadakis. Physics-informed neural networks (pinns) for fluid mechanics: A review. Acta Mechanica Sinica, 37(12):1727–1738, 2021.

[19] Georgios Kissas, Yibo Yang, Eileen Hwuang, Walter R. Witschey, John A. Detre, and Paris Perdikaris. Machine learning in cardiovascular flows modeling: Predicting arterial blood pressure from non-invasive 4D flow mri data using physics-informed neural networks. Computer Methods in Applied Mechanics and Engineering, 358:112623, 2020.

[20] Tianping Chen and Hong Chen. Universal approximation to nonlinear operators by neural networks with arbitrary activation functions and its application to dynamical systems. IEEE Transactions on Neural Networks, 6(4):911–917, 1995.

[21] Lu Lu, Pengzhan Jin, Guofei Pang, Zhongqiang Zhang, and George E. Karniadakis. Learning nonlinear operators via DeepONet based on the universal approximation theorem of operators. Nature Machine Intelligence, 3(3):218–229, 2021.

[22] M. Sherburn, D. G. Chang, and A. J. Engler. Breast duct wall mechanics and viscoelastic characterisation. Biomechanics and Modeling in Mechanobiology, 13:1– 12, 2014.

[23] Lin Jiang, Diana L Alatalo, and Fatemeh Hassanipour. A human milk-mimicking fluid for piv experiments. Experiments in Fluids, 61(10):224, 2020.

[24] Suncica Caníć and Eun Heui Kim. Mathematical analysis of the quasilinear effects in a hyperbolic model of blood flow through compliant axi-symmetric vessels. Mathematical Methods in the Applied Sciences, 26(14):1161–1186, 2003.

[25] Nilza G Ramião, Pedro S Martins, Rita Rynkevic, António A Fernandes, Maria Barroso, and Diana C Santos. Biomechanical properties of breast tissue, a state-of-the-art review. Biomechanics and modeling in mechanobiology, 15(5):1307–1323, 2016.

[26] Malcolm M. Cross. Rheology of non-Newtonian fluids: a new flow equation for pseudoplastic systems. Journal of Colloid Science, 20(5):417–437, 1965.

[27] Diederik P. Kingma and Jimmy Ba. Adam: A method for stochastic optimization, 2017.

[28] Luca Formaggia, Jean-Frédéric Gerbeau, Fabio Nobile, and Alfio Quarteroni. Numerical treatment of defective boundary conditions for the Navier-Stokes equations. SIAM Journal on Numerical Analysis, 40(1):376–401, 2003.

[29] John R. Womersley. Method for the calculation of velocity, rate of flow and viscous drag in arteries when the pressure gradient is known. Journal of Physiology, 127(3):553–563, 1955.

[30] Yann A LeCun, Léon Bottou, Genevieve B Orr, and Klaus-Robert Müller. Efficient backprop. In Neural Networks: Tricks of the Trade, pages 9–48. Springer, 2012.

[31] Zongyi Li, Nikola Kovachki, Chris Choy, Boyi Li, Jean Kossaifi, Shourya Otta, Mohammad Amin Nabian, Maximilian Stadler, Christian Hundt, Kamyar Azizzadenesheli, et al. Geometry-informed neural operator for large-scale 3d pdes. Advances in Neural Information Processing Systems, 36, 2024.

